# Gram-negative bacterial infection increases lung cancer metastasis via Toll-like receptor activation and increased cancer cell proliferation post-tumor adhesion

**DOI:** 10.1101/2021.09.15.460508

**Authors:** Roni F. Rayes, Marnie Goodwin Wilson, Stephen D. Gowing, France Bourdeau, Betty Giannias, Sophie Camilleri-Broet, Pierre-Olivier Fiset, Sara Najmeh, Jonathan D. Spicer, Lorenzo E. Ferri, Jonathan J. Cools-Lartigue

**Author notes:** <u>Corresponding author:</u> Jonathan Cools-Lartigue - MD PhD, Assistant Professor, Department of Surgery, McGill University, QC, Canada. These authors contributed equally to this work (Roni F. Rayes, Marnie Goodwin Wilson).

## Abstract

**Background:** Lung cancer is a leading cause of death partially due to high recurrence rates after surgical resection. Clinical data suggest that post-operative infections may increase the risk of recurrence. Our previous work indicated that increased adhesion of circulating tumors in the context of infection is partially responsible for this phenotype. However, cancer metastasis is a multi-step process, and it is likely that other events following tumor adhesion also play a role.

**Methods:** *In vivo* intrasplenic injection of murine lung cancer cells into wild type (WT) and Toll-like receptor 4 knockout (TLR4^-/-^) mice followed by cecal-ligation and puncture (CLP) as a model of post-operative infection or sham surgery were used. H&E staining and immunohistochemistry analysis of Ki67+ cells in the livers of those mice were performed. *In vitro* proliferation assays were performed on human lung cancer cells using combinations of TLR blockade.

**Results:** We found a 5-fold increase in hepatic metastases in WT CLP mice compared to WT sham mice. TLR4^-/-^ CLP mice had a significant decreased tumor burden compared to WT CLP mice. This indicated an important mechanistic role for the TLR4-initiated host response to gram negative infection post-tumor cell adhesion. By analyzing the livers of those mice, we observed an increase in proliferation of tumor micrometastases *in vivo* in WT CLP mice as compared to WT sham mice. Here again, CLP TLR4 ^-/-^ mice had significantly fewer replicating micrometastases than CLP WT mice. Indeed, we found that direct stimulation of lung cancer cells with heat-inactivated *E*.*Coli* resulted in increased proliferation of tumor growth *in vitro*. These effects were partially abrogated by tumor TLR4 blockade; combined TLR2, 4 and 5 blockades led to a more prominent decrease. Conditioned media from bronchoalveolar epithelial cells treated with lipopolysaccharide lead to increased lung cancer proliferation; these changes were reversed with TLR blockade, indicating that the host response to infection is TLR mediated.

**Conclusions:** Overall, these results imply a more complex mechanistic role of post-operative infection in metastasis. From a clinical standpoint, this evidence strengthens the case for the use of TLR blockade as a potential therapeutic target in the prevention of metastasis.

## Introduction

Lung cancer is the leading cause of cancer-related mortality world-wide. Prognosis improves with early detection due to the potential for curative surgical resection, but therapeutic success in the long-term is hampered by high rate of metastatic recurrence after surgery. Two mechanisms are proposed to explain the emergence of metastases after surgical resection: (i) tumor manipulation during surgery can cause introduction of circulating tumour cells (CTCs) into the bloodstream or lymphatic system (1), leading to their adhesion to secondary sites; (ii) patients undergoing cancer surgery already harbor radiologically undetectable micro-metastases, which can serve as reservoirs for future metastases (2).

Research from our group suggest that post-operative infections following tumour resection, a frequent complication of surgery, may contribute to these high recurrence rates (3). Indeed, 25-40% of patients undergoing lung cancer resection will have some type of postoperative complication, the majority of which are infectious. Gram-negative organisms account for a significant proportion of post-operative infections; in particular, gram-negative pneumonia is a well-known complication following lung cancer resection (4). The pro-inflammatory state caused by gram-negative infection is largely due to lipopolysaccharide (LPS). LPS is recognized by Toll-like receptor 4 (TLR4), a highly conserved pattern recognition receptor involved in the initiation of the immune response. Similar responses to gram-negative infection are also induced by other TLRs, including TLR2 and TLR5 (5). We have previously shown that TLR4 activation by gram-negative pathogens increases solid cancer metastases in murine models by increased adhesion of CTCs (6). Though increased cancer cell adhesion to metastatic sites provides an elegant explanation if one assumes that this phenotype is entirely related to CTCs, it is less likely that adhesion has such a significant role in the large proportion of patients already harboring radiologically undetectable distant metastases. Instead, other mechanisms are involved since the pathogenesis of metastasis includes both pre- and post-adhesive events (7).

This paper investigates the role that bacterial infection plays in promoting the formation of lung cancer metastasis by examining the mechanisms of tumor spread post-adhesion. Previous work has indicated that both host- and tumor-derived factors play a role in the increased tumor burden following post-operative infection. Indeed, many tumor cells are known to express functional TLRs which allow the to recognize and initiate signaling cascades in response to LPS. In addition, gram-negative sepsis can induce significant inflammatory changes in the tumor immune microenvironment (TIME) that can lead to increased cancer cell replication (8). Finally, the inflammatory cascade triggered by TLRs, particularly TLR4, may be a therapeutic target to reverse this phenomenon. In fact, blockade of TLR4 has previously been investigated and shown to be safe in the treatment of gram-negative sepsis (9). If these events can be abrogated via blockade of the inciting factors this work could have significant implications for patients undergoing surgical resection of cancer.

## Materials and Methods

### Animals

7-9 weeks old male C57BL/6 (Charles River, Saint Constant, QC) and TLR4^-/-^ (from Dr. Qureshi (McGill)) mice were used in all experiments and are approved by the McGill Animal Care Committee and conducted in accordance with institutional guidelines. In this study, only male mice were used to reduce the effect of sex hormones (namely estrogen). A sample size of 7 mice per cohort will be used to achieve a power of 80% and a level of significance of 5% (2-sided), for detecting a true difference in means of 25%, assuming a pooled standard deviation of 15 units.

### Anesthetics and analgesics for animal work

To minimize the effects of the proinflammatory mediator phenomenon around the cancer cells and for a better control of the pain, Buprenorphine slow release was injected (1mg/kg good for 72 hours) subcutaneously 30 minutes prior to the start of the surgery. Buprenorphine 0.1 mg/ml diluted in 1mL of NaCl was used after 72 hours. Animals were anesthetized with Isoflurane 4% (inhalant) and maintained under isoflurane at 2%. Ophthalmic ointment was applied to prevent dryness and damage to the cornea. Body temperature was maintained using heating disc or warming pad. A mixture of Lidocaine/bupivacaine was applied on the wound following CLP and intrasplenic injection just before closure. A 1:1 mixture of lidocaine HCl 2% (20mg/ml) injectable solution and bupivacaine HCl 0.50% (5mg/ml) injectable solution was used. Animal were placed gently on a thermal blanket, monitored until they awaken from anesthesia and then returned to their cages. The cage was then placed in a mouse incubator before returning to the animal room.

### Animal monitoring parameters

Mice were monitored 2-3 times a day for the first week and daily for the second week after CLP and intrasplenic injection. They were injected with buprenorphine or NaCl as needed until endpoint between day 10-14. Mice have food at the bottom of their cage as well as gel food or Ensure. Their anus are also gently cleaned with warm water and ointment or zinc oxide cream (Sudocream) is applied. Should any of the following clinical endpoints manifest ie. signs of distress such as failure to right themselves, severe pain on touch or agonal breathing, the animals were euthanized. We have ∼5-10% mortality due to septicemia.

### Reagents

LPS derived from *Escherichia coli* (*E. coli*) was from Sigma-Aldrich Canada (Oakville, ON). *E. coli* were heat-inactivated at 95°C, 10min and cooled to room temperature (RT) prior to use. Eritoran (TLR4 antagonist) was from Eisai (Mississauga, ON); BIRB0796 (p38 MAPK inhibitor) was from Professor Cohen (University of Dundee). PD184352 (MEK1/2 inhibitor) was from US Biological (Swampscott, MA). PI103 (PI3K inhibitor) was from Cayman Chemical (Ann Arbor, MI)

### Cell culture

Murine Lewis lung carcinoma (LLC) subline H59, stably expressing GFP (H59-GFP), from Dr. Brodt (McGill), were maintained in RPMI, 10% FBS, 1% glutamine, 1% penicillin-streptomycin (pen/strep). Human lung adenocarcinoma cell line A549, obtained from the American Type Culture Collection (ATCC), was maintained in DMEM/F12, 10% FBS, 1% pen/strep. Human lung bronchoepithelial cell line BEAS-2B, obtained from ATCC, was maintained in RPMI with 10% FBS, 1% glutamine, 1% pen/strep. All cell culture reagents are from Wisent Bioproducts (St-Bruno, QC). All experiments were performed with mycoplasma-free cells at 37°C, 5% CO_2_. Human cell lines have been authenticated using STR profiling within the last three years.

Prior to experiments, H59-GFP and A549 cells were pre-incubated for 24hrs in serum-free (SF) media. Eritoran, PD184352, BIRB0796, and PI103 were all used at 100 nM and were added 1hr prior to incubation with 100 ng/mL LPS or 10^8^ colony-forming units/mL of heat-inactivated *E. coli* and remained in media during stimulation.

BEAS-2B cells were stimulated with 100 ng/mL LPS for 2hrs, starved 24hrs in SF-DMEM, following which conditioned media was collected and supplemented with 10% FBS. Eritoran was added to SF-media 30 min prior to LPS stimulation and remained in media during stimulation.

### Antibodies

Neutralizing recombinant IgA2 monoclonal antibodies to human TLR5 and TLR2 and their human IgA2 isotype control were used at 0.1 μg/mL. Neutralizing IgG2a rat monoclonal antibodies to murine TLR5 and TLR2 were used at 0.1 and 1 μg/mL, respectively. The rat IgG2a isotype control was used at the same concentrations. All TLR antibodies were from InvivoGen.

Fluorescein-conjugated monoclonal rat anti-mouse TLR2 IgG2a (R&D systems (Minneapolis, MN)) was used at 50 μg/mL. Fluorescein-conjugated monoclonal mouse anti-mouse TLR5 IgG2a and fluorescein-conjugated monoclonal rat anti-mouse TLR4 IgG2b (both from Novus Biologicals (Oakville, ON)) were used at 500 μg/mL. Fluorescein-conjugated monoclonal mouse IgG2a, fluorescein-conjugated monoclonal rat IgG2a and fluorescein-conjugated monoclonal rat IgG2b were used at identical concentrations as isotype controls (all from R&D systems). All antibodies were incubated for 1hr with cells in 0.1M Tris-HCl, 10% FBS in PBS at 4°C.

### Staining with CellTracker Red

H59-GFP cells were stained with CellTracker Red according to manufacturer’s instructions and analyzed by flow cytometry (FACScan, BD Biosciences, Mississauga, ON).

### Cell proliferation assay

5,000 cells in SF-media were seeded into 96-well plates, incubated for 24hrs, treated with *E. coli* ± Eritoran or conditioned media from BEAS-2B cells, and proliferation assessed by MTT assay according to manufacturer’s instructions (R&D Systems).

### Liver metastasis assay

C57BL/6 male mice received an intrasplenic (i.s.) injection of 3×10^4^ cells followed by splenectomy. 24hrs later, peritonitis was induced by cecal-ligation-puncture (CLP) as previously described (10). Sham animals had their cecums exteriorized without CLP. Mice were sacrificed after 2 weeks and their livers were harvested, visible metastases counted, and representative images captured.

### Immunohistochemistry

Livers were formalin-fixed, paraffin-embedded, sectioned, blocked and stained with monoclonal rabbit anti-Ki-67 antibody (1:400; Cell Signaling, Boston, MA), secondary goat anti-rabbit antibody (1:500; Jackson Lab, Bar Harbor, ME) and DAB substrate kit (DAKO, Burlington, ON). Sections were counterstained with Hematoxylin, visualized and photographed using an inverted light microscope with 10X, 20X and 100X objectives (Nikon TE300) and a digital SLR camera system (Nikon D90).

### Statistical analysis

Data were analyzed using T-test (for normally distributed data) or Mann-Whitney test (for not-normally distributed data) using GraphPad (Prism). Normal distribution was assessed using the Kolmogorov-Smirnov test. P < 0.05 will be considered statistically significant. WT and KD animals were randomized in the CLP and sham groups and all analysis were done blinded. Animals were mixed withing a cage and fighting animals were separated to avoid inflammation.

## Results

### Post-operative infection increases lung cancer metastasis via TLR4 activation

Previous work has shown that in mice with active infection, i.s. injection of tumor cells results in increased liver metastatic burden (3). To better replicate the clinical model of post-operative infection and to determine whether post-adhesive events were contributory to this phenotype, we intrasplenically injected C57BL/6 wild type (WT) mice with H59-GFP LLC cells, followed by CLP or sham surgery 24hrs after, to induce gram-negative bacterial sepsis after tumor cell injection. WT mice that underwent CLP demonstrated a significant 5-fold increase in hepatic metastases compared to those that underwent sham surgery (p=0.0005; **Fig.1A,B**). To investigate whether TLR4 activation contributed to this increase in H59-GFP hepatic metastases, the experiment was repeated with TLR4^-/-^ mice. In TLR4^-/-^ mice, a smaller and less significant 2-fold increase in hepatic metastases was observed in mice that underwent CLP compared to those that received sham surgery (p=0.0082; **Fig.1A,B**). A statistically significant difference in hepatic metastases was also observed between WT and TLR4^-/-^ mice that underwent CLP (p=0.0006; **Fig.1A,B**) but not between the sham groups (p=0.2168; **Fig.1A,B)**.

**Fig. 1.**
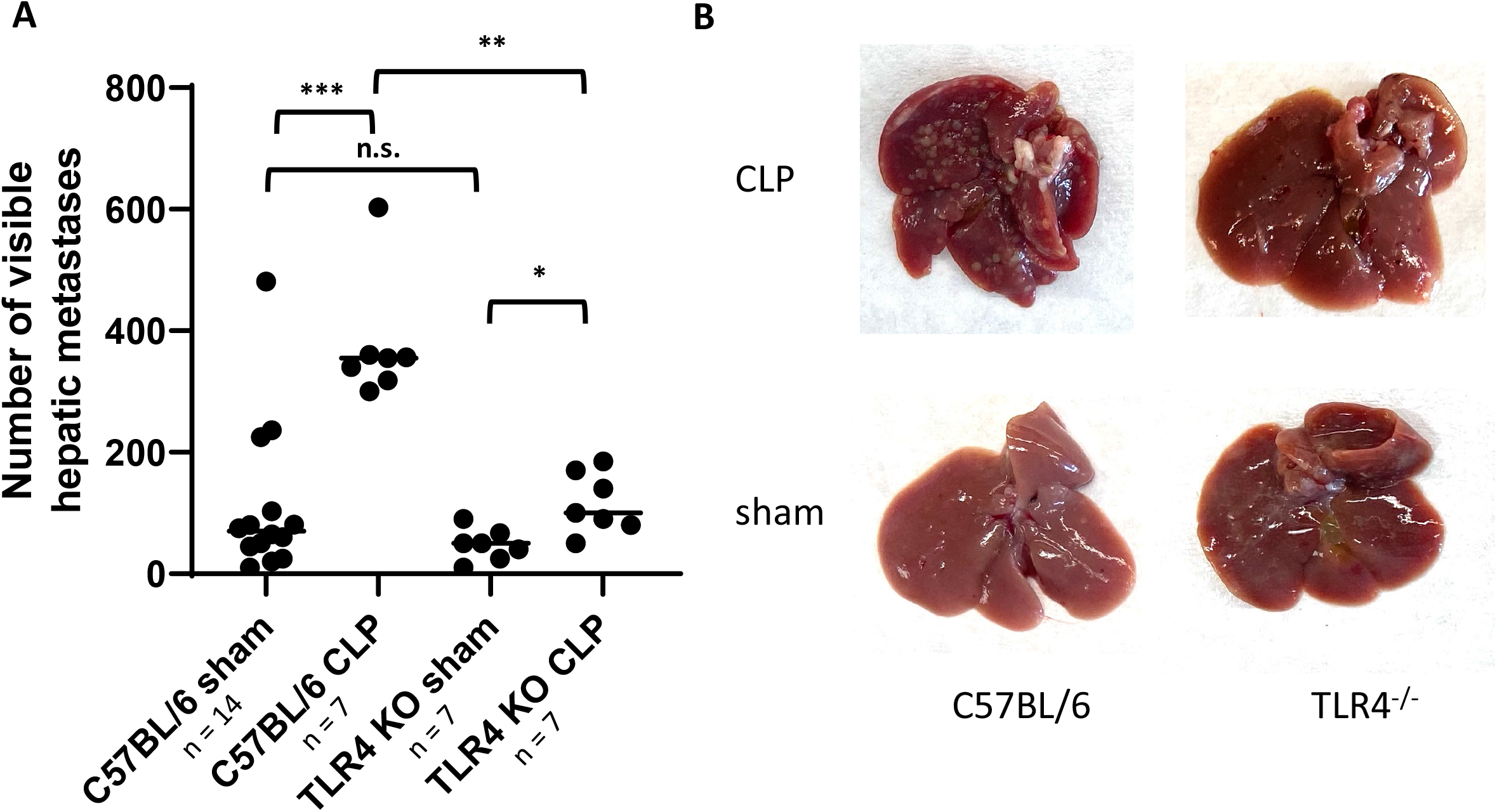
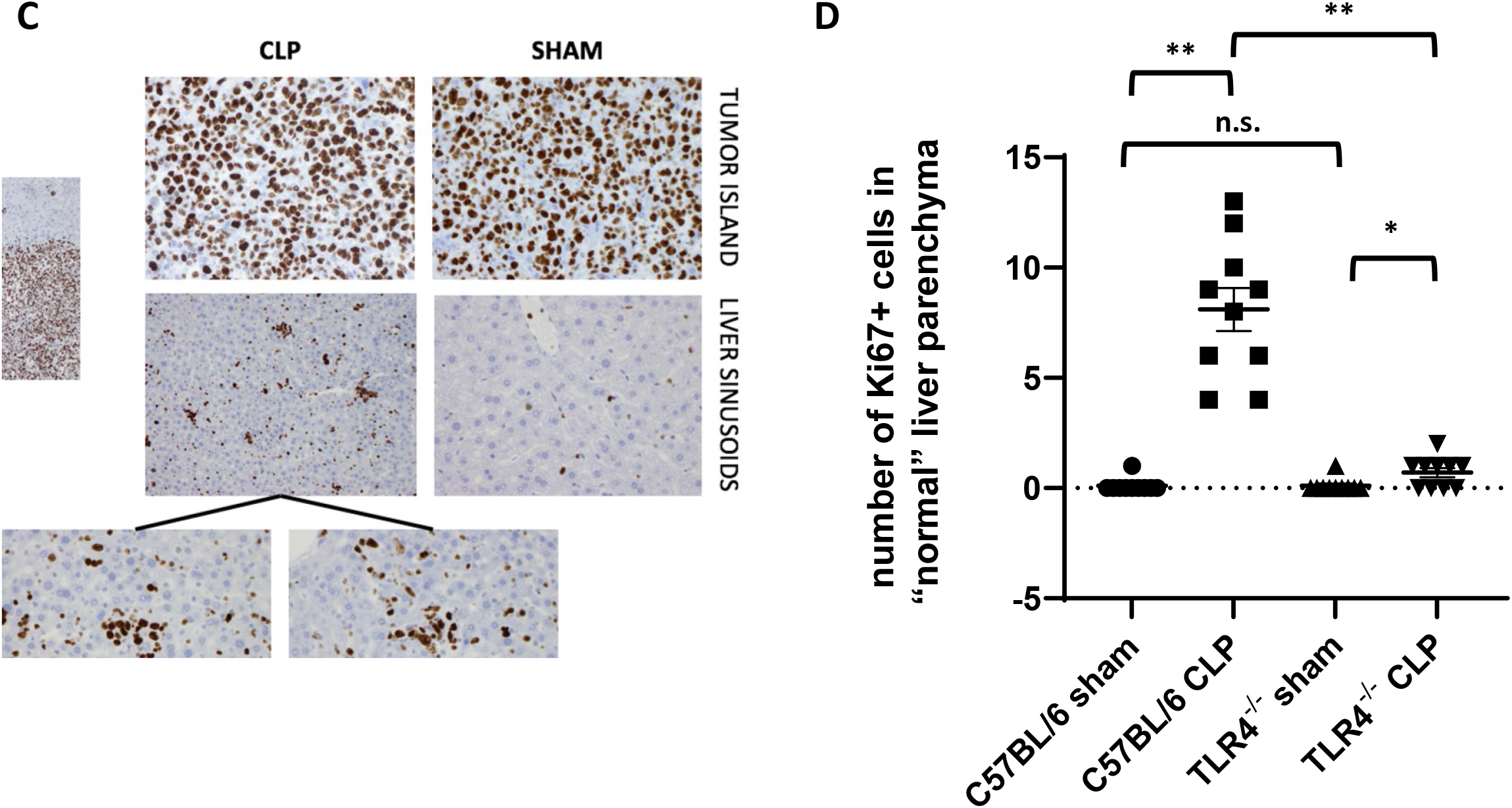
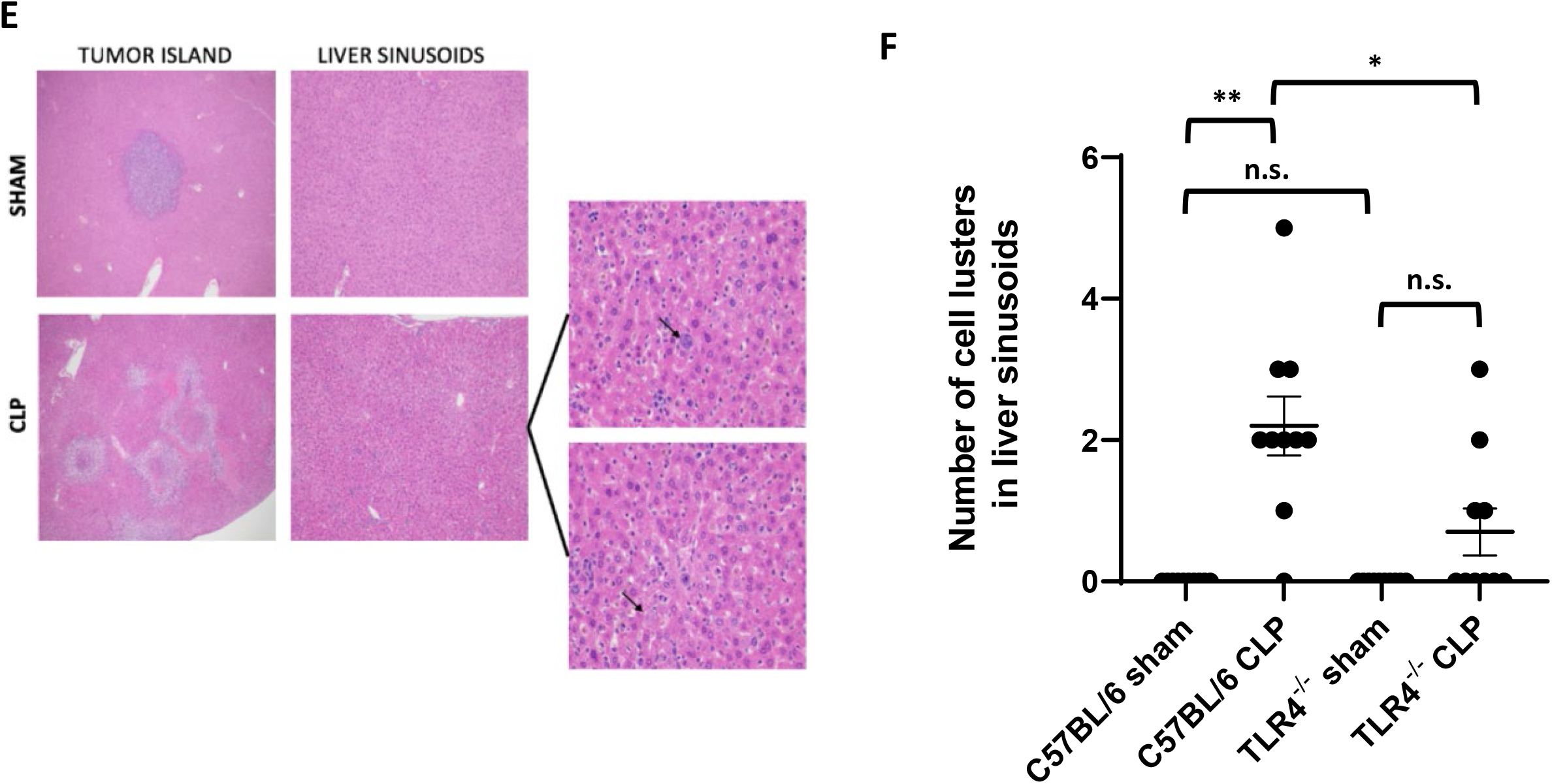
Murine H59-GFP liver metastasis is augmented *in vivo* by CLP and resultant gram-negative sepsis 24 hours post-tumor cell injection. **A**. Dot plot of number of gross hepatic metastases in WT and TLR4^-/-^ mice injected with 30,000 H59-GFP cells followed by CLP or sham surgery 24hrs later and sacrificed 2 weeks after tumor cell injection with means and standard deviations. ^*^ p = 0.0082; ^**^ p = 0.0006; ^***^ p = 0.0005. **B**. Representative images of the livers from the 4 groups of mice in (A). **C**. Representative images of Ki67 staining in tumor island and liver sinusoids of livers from sham and CLP mice two weeks following tumor injection of H59-GFP cells followed by CLP after 24hrs using a 20X microscope objective. **D**. Dot plot of number of ki67 tumor islands seen in 10 random fields (± SEM) from the 4 groups of mice in (A). ^*^ p = 0.0203; ^**^ p < 0.0001. **E**. Representative H&E images of the livers of the 4 groups of mice in (A) at 20X magnification and inset of micrometastases of the micrometastases seen in the CLP group at 40X magnification. **F**. Dot plot of number of micrometastases seen in 10 random fields (± SEM) from the 4 groups of mice in (A). ^*^ p = 0.117; ^**^ p < 0.0001. For all panels, n=7-14 mice/group, n.s.=not significant.

### Post-operative infection increases proliferation of tumor cell micrometastases in vivo via TLR4 activation

The collected livers were formalin-fixed and paraffin-embedded and stained with ki67 and assessed by immunohistochemistry (IHC). As expected, in all groups, tumor metastases uniformly showed 90-100% Ki-67 positivity (Ki67+) compared to background normal liver parenchyma **(Fig.1C)**. Importantly, WT mice that underwent CLP had significantly increased levels of Ki67 in macroscopically normal liver parenchyma as compared to mice that underwent sham surgery, indicating an increase in tumor cell replication in the context of gram-negative sepsis (p< 0.0001; **Fig.1C,D**). This was significant lower in CLP TLR4^-/-^ mice compared CLP WT mice (p< 0.0001; **Fig.1D**). The increase in ki67+ cells in WT CLP mice corresponded to a significant increase in microscopic clusters of cells within the liver sinusoids of those mice as compared to sham (p<0.0001; **Fig.1E,F**), consistent with replicating tumor micrometastases within their liver sinusoids. Here again, the CLP TLR4^-/-^ mice had significantly fewer replicating micrometastases than the CLP WT mice (p=0.0117; **Fig.1F**).

We then investigated whether increased cancer cell proliferation is implicated in the mechanism of increased tumor burden in WT mice undergoing CLP, by testing both (i) the effect of direct stimulation of the cancer cells with LPS and (ii) the effect of the TIME, as modified by the presence of gram-negative sepsis.

### TLR activation increases proliferation of murine LLC cells in vitro and can be abrogated by TLR blockade

To test this hypothesis, we first focused on the effects of TLR4 stimulation on the cancer cell directly. H59-GFP cells were plated and stimulated with heat-inactivated *E. coli*, with or without Eritoran, a small-molecule TLR4 inhibitor. At 24, 48- and 72-hours post-stimulation, we observed increase confluence in cells treated with *E. coli* as compared to untreated cells, although this difference did not become statistically significant until 48 hours post-treatment **(Fig.2A,B)**. Moreover, at all time points, partial abrogation of this effect was seen in samples treated with Eritoran as compared stimulated with *E. coli*, although again this difference did not become statistically significant until 72 hours post-treatment (p<0.05; **Fig.2A,B)**.

**Fig. 2.**
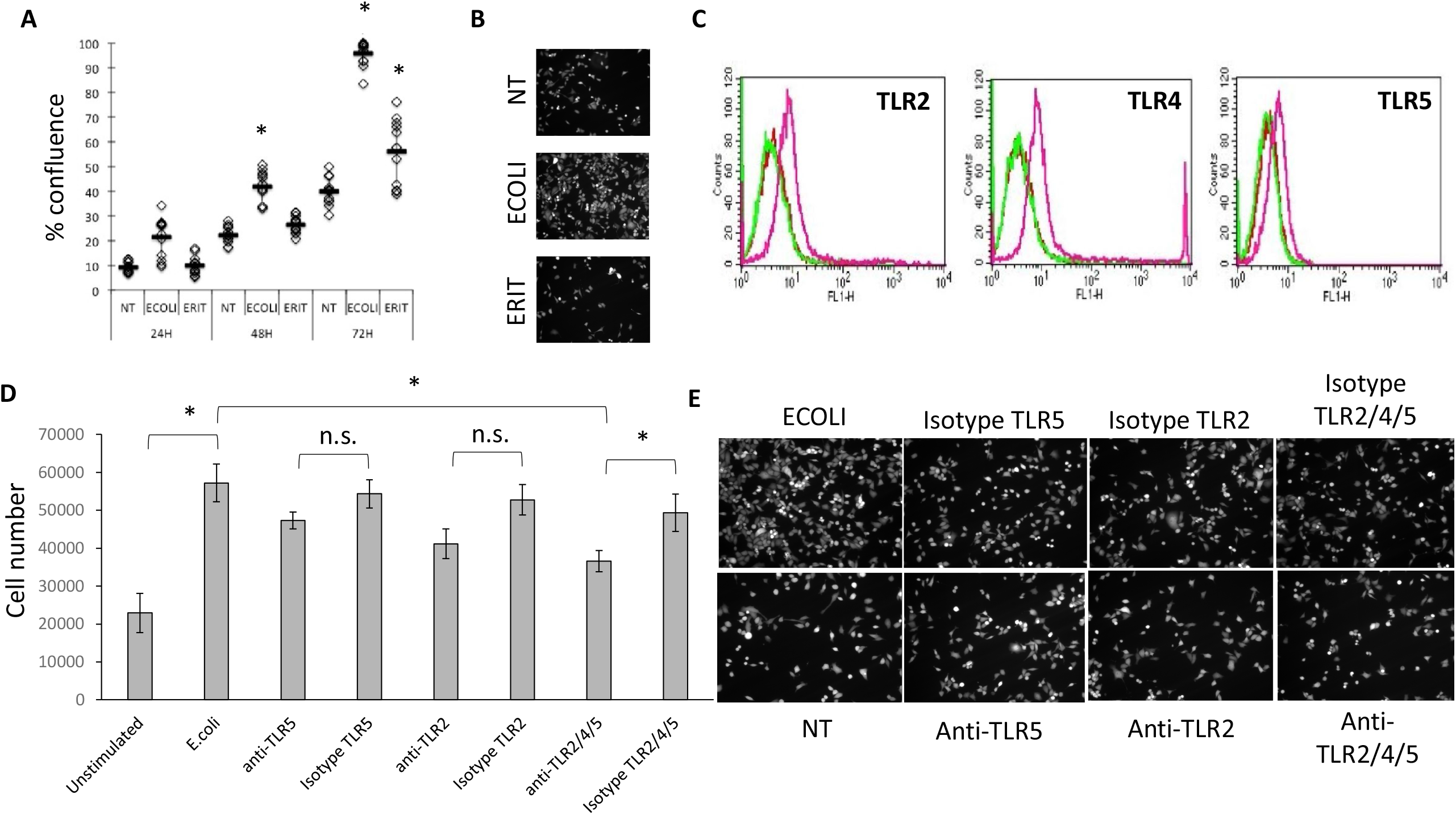
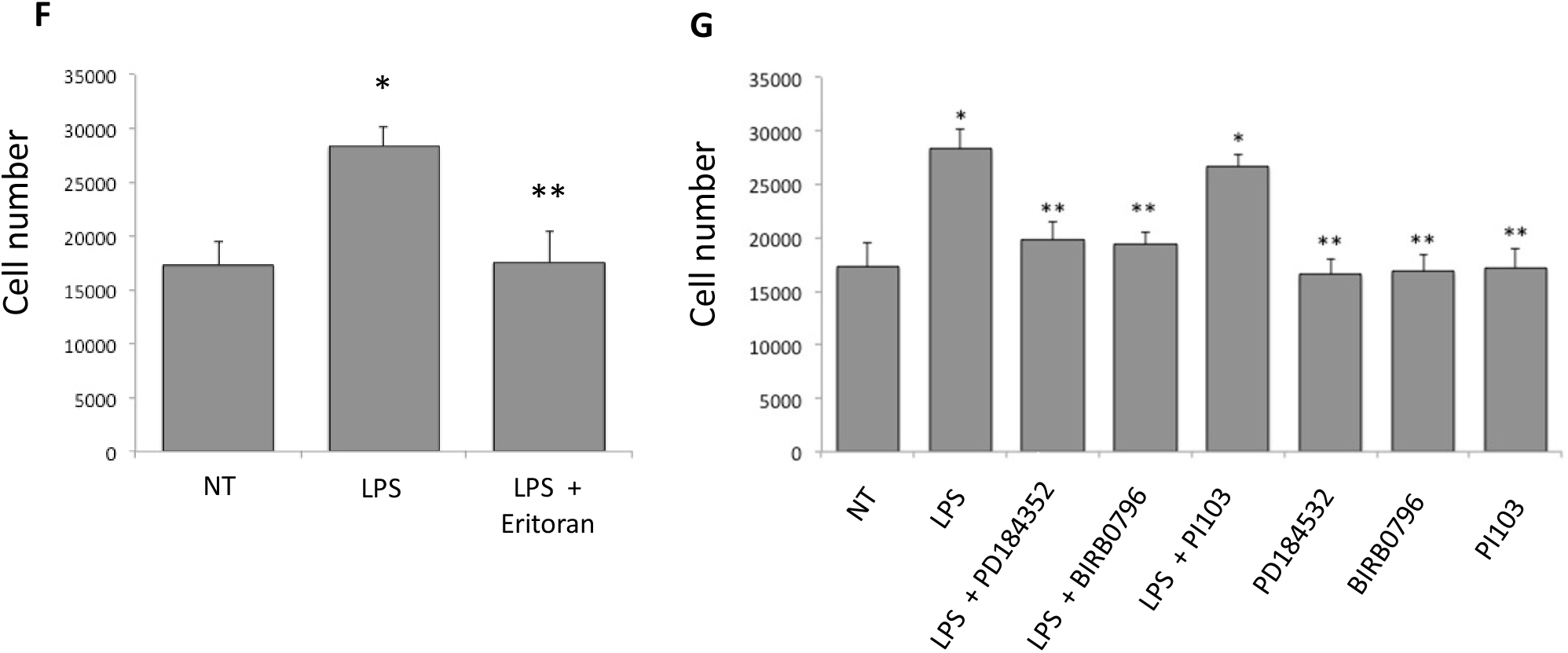
Lewis Lung Carcinoma cell proliferation is increased by heat-inactivated *E. Coli* and is partially abrogated by TLR blockade. **A**. Dot plot number of % confluence of GFP+ cells (average ± SEM) from cultures of H59-GFP cells stimulated with either heat inactivated *E. coli* (ECOLI) or a combination of heat inactivated *E. coli* and Eritoran (ERIT) at 24, 48 and 72 hrs post-stimulation. Non-treated (NT) cells were used as controls. n=9, ^*^p<0.05. **B**. Representative images of panel A. **C**. Cell surface expression of H59-GFP by flow cytometry for TLR2, TLR4 and TLR5. For all 3 panels, non-treated cells are in green, isotype control in red and antibody against TLR in pink. 10,000 cells were counted for each sample. **D**. Bar graph of average (± SEM) number of H59 cells following stimulation with heat-inactivated *E. Coli* in the presence or absence of blockade of TLR2, TLR5 and TLR2, 4 and 5 combination along with their isotype controls. n≥3, ^*^p<0.05. **E**. Representative images of panel D. **F**. Bar graph of MTT assay 48 hours post-stimulation with conditioned media from untreated BEAS-2B cells (NT), BEAS-2B cells stimulated with LPS (LPS), BEAS-2B cells stimulated with LPS following treatment with small molecule inhibitor Eritoran (LPS + Eritoran). n≥3, n.s.=not significant, ^*^p<0.05 increase from NT, ^**^p<0.05 decrease from LPS. **G**. Bar graph of MTT assay 48 hours post-stimulation with conditioned media from untreated BEAS-2B cells (NT), BEAS-2B cells stimulated with LPS (LPS) and BEAS-2B cells stimulated with LPS following treatment with small molecule inhibitors PD184352, BIRB0796 and PI103 (LPS + “inhibitor”) or inhibitor alone. Data are presented as mean ± standard deviation. n≥3, ^*^p<0.05 increase NT and ^**^p<0.05 decrease from LPS

Although significant decreases in cell replication were observed in cells pre-treated with Eritoran, growth of cells stimulated with heat inactivated *E. coli* consistently remained above growth of non-treated cells **(Fig.2A,B)**. Since the heat inactivated *E. coli* strain we use in this study is known to express Flagellin, a pathogen associated molecular pattern detected by TLR5 and that TLR2 is known to initiate responses to peptidoglycan, a component of gram negative bacteria, we hypothesized that both TLR2 and TLR5 might be also playing a role in the proliferation of lung cancer cell lines following sepsis. Using flow cytometry, we first show that H59-GFP cells express TLR2, TLR4 and TLR5 **(Fig.2C)**. Blockade of TLR2 and TLR5 individually resulted in a slight abrogation of H59 cell proliferation; however, none of these effects were statistically significant (p>0.05; **Fig.2D,E**). Co-blockade of all 3 receptors resulted in significant decrease of H59 cell proliferation compared to cells stimulated but not TLR blocked (p<0.05; **Fig.2D,E**). But once again, it did not represent a complete return to baseline rates of cell proliferation in non-treated cells.

### A549 cancer cell proliferation is augmented by conditioned media collected from BEAS-2B cells stimulated with LPS through ERK and p38 MAP Kinase dependent pathways

After investigation of the effect of direct stimulation of tumor cells with heat inactivated *E. coli*, we turned our attention to the effect of the host immune response on tumor cell proliferation. We observed, that A549 cells treated with conditioned media from BEAS-2B cells stimulated with LPS, showed increased replication compared to cells treated with media from non-treated BEAS-2B cells or BEAS-2B cells pre-treated with Eritoran prior to LPS stimulation (p<0.05; **Fig.2F**). In this case, the increased replication was completely abrogated by TLR4 blockade with Eritoran (**Fig.2F**).

Given that stimulation of BEAS-2B cells with LPS enhanced cell replication, we sought to determine whether this mechanism was dependent on activation of downstream cascades of TLR4. When BEAS-2B cells were pre-treated with either PD184352 or BIRB0796, prior to LPS stimulation, A549 cell replication returned to levels of non-treated cells (p<0.05; **Fig.2G**). This was not the case when using conditioned media from BEAS-2B cells pre-treated with PI103 prior to LPS stimulation (p>0.05; **Fig.2G**).

## Discussion

We have previously shown that the initiation of the innate immune response through the stimulation of TLR4 results in increased tumor cell adhesion, offering a partial explanation for this phenomenon in patients with CTCs (6, 11). However, this mechanism does not account for the significant proportion of patients thought to have radiologically undetectable metastatic disease prior to surgery. These patients are at high risk for early recurrence, and it seems likely that post-operative infection could have deleterious effects with regards to recurrence rate on this population as well.

In this work, we show that stimulation of various TLRs expressed on tumor cells, as well as the stimulation of TLR4 in the host, results in increased replication of lung cancer cells, both *in vitro* and *in vivo*, in a highly physiologic model of gram-negative infection. Moreover, we show that this result is at least partially abrogated in TLR4^-/-^ mice, indicating a potential role for therapeutic blockade of TLR4 to inhibit this phenomenon. *In vitro*, the small molecule inhibitor Eritoran, which targets TLR4, shows significant potential in the abrogation of the host response and resultant increased tumor cell proliferation, while blockade of TLR4, TLR2 and TLR5 all showed a potential effect in decreasing proliferation of tumor cells directly stimulated with heat inactivated E. Coli. These results make sense in that the NF-kB pathway, which is initiated by TLR activation, has been previously shown to have a role in cell mitosis and has been implicated in oncogenesis (12, 13). In addition, the PI3K cascade, which is one of the initiators of the NF-kB-mediated response to infection, has been targeted in the past for cancer-cell directed small molecule inhibitors of replication (6).

With regards to the role of the host response to infection, which is thought to be the major driver of the tumor cell microenvironment, both our *in vitro* and *in vivo* experiments have demonstrated that TLR4 activation may be one of the key components in driving increased tumor cell replication. Importantly, blockade of this pathway has already been investigated and has been shown to be safe in patients with gram-negative sepsis outside of the context of cancer (9). While blockade of TLR4 has not been shown to be beneficial in the gram-negative sepsis population, these results indicate a potentially important role for TLR4 blockade as a therapeutic modality to decrease post-operative recurrence of lung cancer.

## Conclusions

Recurrence of lung cancer after surgical resection with curative intent is a significant problem, both in terms of public health and with regards to the individual patient. Epidemiologically, post-operative infection increases risk of early metastatic recurrence in lung cancer patients. Currently, no therapeutic options are available that target this phenomenon. Our data here demonstrate a more complex mechanistic role of post-operative infection in metastasis. From a clinical standpoint, this evidence strengthens the case for the use of TLR blockade as a potential therapeutic target in the prevention of metastasis.

## Acknowledgments

We would like to thank the histopathology platform of the RI-MUHC for their help with the immunohistochemistry.

## List of abbreviations

TLR4: Toll-like receptor 4
CTCs: circulating tumour cells
LPS: lipopolysaccharide
TIME: tumor immune microenvironment
E. coli: Escherichia coli
LLC: Lewis lung carcinoma
ATCC: American Type Culture Collection
SF: serum-free
i.s.: intrasplenic
CLP: cecal-ligation-puncture
WT: wild type
IHC: immunohistochemistry

## Declarations

### Ethics approval and consent to participate

the animal experiments were approved by the McGill Animal Care Committee and conducted in accordance with institutional guidelines.

### Consent for publication

not applicable.

### Availability of data and material

The datasets used and/or analysed during the current study are available from the corresponding author on reasonable request.

### Competing interests

The authors declare that they have no competing interests.

### Funding

Canadian Institutes of Health Research; Grant number: MOP-133567.

### Authors’ contributions

- Conceptualization: MGW, SDG, SN, JDS, LEF, JJCL
- Experiments: RFR, MGW, SDG, FB, BG, SCB, POF, SN
- Data analysis: RFR, MGW, SDG, FB, BG, SCB, POF, SN
- Writing: RFR, MGW, SDG, FB, BG, SCB, POF
- Supervision: SN, JDS, LEF, JJCL.

## Notes

### Competing Interest Statement

The authors have declared no competing interest.

